# Improvement of the threespine stickleback (*Gasterosteus aculeatus*) genome using a Hi-C-based Proximity-Guided Assembly method

**DOI:** 10.1101/068528

**Authors:** Catherine L. Peichel, Shawn T. Sullivan, Ivan Liachko, Michael A. White

## Abstract

Scaffolding genomes into complete chromosome assemblies remains challenging even with the rapidly increasing sequence coverage generated by current next-generation sequence technologies. Even with scaffolding information, many genome assemblies remain incomplete. The genome of the threespine stickleback (*Gasterosteus aculeatus)*, a fish model system in evolutionary genetics and genomics, is not completely assembled despite scaffolding with high-density linkage maps. Here, we first test the ability of a Hi-C based proximity guided assembly to perform a *de novo* genome assembly from relatively short contigs. Using Hi-C based proximity guided assembly, we generated complete chromosome assemblies from 50 kb contigs. We found that 98.99% of contigs were correctly assigned to linkage groups, with ordering nearly identical to the previous genome assembly. Using available BAC end sequences, we provide evidence that some of the few discrepancies between the Hi-C assembly and the existing assembly are due to structural variation between the populations used for the two assemblies or errors in the existing assembly. This Hi-C assembly also allowed us to improve the existing assembly, assigning over 60% (13.35 Mb) of the previously unassigned (∼21.7 Mb) contigs to linkage groups. Together, our results highlight the potential of the Hi-C based proximity guided assembly method to be used in combination with short read data to perform relatively inexpensive *de novo* genome assemblies. This approach will be particularly useful in organisms in which it is difficult to perform linkage mapping or to obtain high molecular weight DNA required for other scaffolding methods.

## Introduction

While short-read genome sequencing has become a staple in genetic research, scaffolding complete eukaryotic genomes from fragmented assemblies remains a remarkably difficult task as most modern scaffolding techniques utilize purified high-molecular weight DNA as the source of contiguity information (Putnam et al. 2016; Teague et al. 2010; Das et al. 2010). Purification results in broken DNA molecules and loss of long-range intra-chromosomal genetic contiguity information typically yielding incomplete scaffolds. One traditional method for retaining chromosome-scale contiguity is to use genetic crosses to establish maps of relative linkage distances between sequences. However, genetic mapping is very laborious, cannot be applied to many organisms, and often falls short of scaffolding all contigs due to low resolution from a limited number of crossovers (reviewed in Fierst 2015). Chromosome conformation capture techniques such as Hi-C (Lieberman-Aiden et al. 2009) retain ultra-long-range genomic contiguity information through *in vivo* crosslinking of chromatin and subsequent sequencing of proximal pairs of sequences. The rate of Hi-C interaction decreases rapidly with increasing genomic distance between pairs of loci. Taking advantage of this relationship between inter-sequence distance and proximity interaction allows the construction of chromosome-scale genome scaffolds (Burton et al. 2013; Marie-Nelly et al. 2014; Kaplan and Dekker 2013).

Here, we used the Hi-C-based Proximity-Guided Assembly method to assemble the genome of the threespine stickleback (*Gasterosteus aculeatus*). This small, teleost fish is a widely used model system in diverse fields, including ecology, evolution, behavior, physiology and toxicology (Wooton 1976; Bell and Foster 1994; Östlund-Nilsson et al. 2007). Sticklebacks are well known for the extensive morphological, behavioral and physiological variation present in freshwater populations that have evolved since the retreat of the glaciers across the Northern hemisphere in the past 15,000 years (Bell and Foster 1994; Hendry et al. 2013). Recent research has led to identification of the genetic and genomic basis of this phenotypic diversity, providing new insights into the genetic basis of adaptation (Peichel and Marques in press). To facilitate research in this model system, a high quality genome assembly for *G. aculeatus* was generated by Sanger sequencing of plasmid, fosmid, and bacterial artificial chromosome (BAC) genomic libraries made from a single female from Bear Paw Lake, Alaska. Scaffolds were anchored to the 21 stickleback chromosomes or linkage groups (LG) using genetic linkage mapping. The original genome assembly comprised 400.7 Mb of scaffolds anchored to linkage groups, with an additional 60.7 Mb of assembled scaffolds not anchored to linkage groups (Jones et al. 2012). Two further revisions to the genome assembly have used genetic linkage mapping in three additional crosses to assign some of these unanchored scaffolds to linkage groups and to correct errors in the original assembly (Roesti et al. 2013; Glazer et al. 2015). The most recent genome assembly comprises 436.6 Mb, with 26.7 Mb remaining unassigned to linkage groups (Glazer et al. 2015). Here, we took advantage of the existence of a high quality assembly for *G. aculeatus* to test the performance of Hi-C in generating a *de novo* assembly. Additionally, we used Hi-C to further improve the existing *G. aculeatus* genome assembly.

## Results

### Hi-C/PGA rescaffolding of the G. aculeatus genome assembly

To investigate how well Hi-C-based Proximity-Guided Assembly (PGA, provided by Phase Genomics, Seattle, WA) can assemble a genome composed of small contigs, we split the revised *G. aculeatus* genome assembly (Glazer et al. 2015) into contiguous 50 kb bins and used a proximity-guided assembly to rescaffold the contigs together. PGA reconstructed a highly accurate genome assembly, with 8,657 of the 8,745 (98.99%) of the 50 kb contigs correctly assigned to one of the 21 linkage groups in the *G. aculeatus* genome during clustering (7 contigs were incorrectly assigned to an alternate linkage group and 81 contigs were not assigned to a linkage group) (Figure 1). Among the linkage groups, nearly all the contigs (8,479 contigs, or 97.94% of the 8,657 contigs in linkage groups) had an identical ordering to the revised *G. aculeatus* reference assembly (Figure 2; Figure S1). The 178 contigs (8.9 Mb of the 436.6 Mb total genome length) that were ordered incorrectly within linkage groups may represent errors in the PGA scaffolding, assembly errors in the reference genome, or could reflect structural variation between the Paxton Lake (British Columbia) benthic population used for the PGA scaffolding and the Bear Paw Lake (Alaska) population used for the reference assembly.

**Figure 1.**
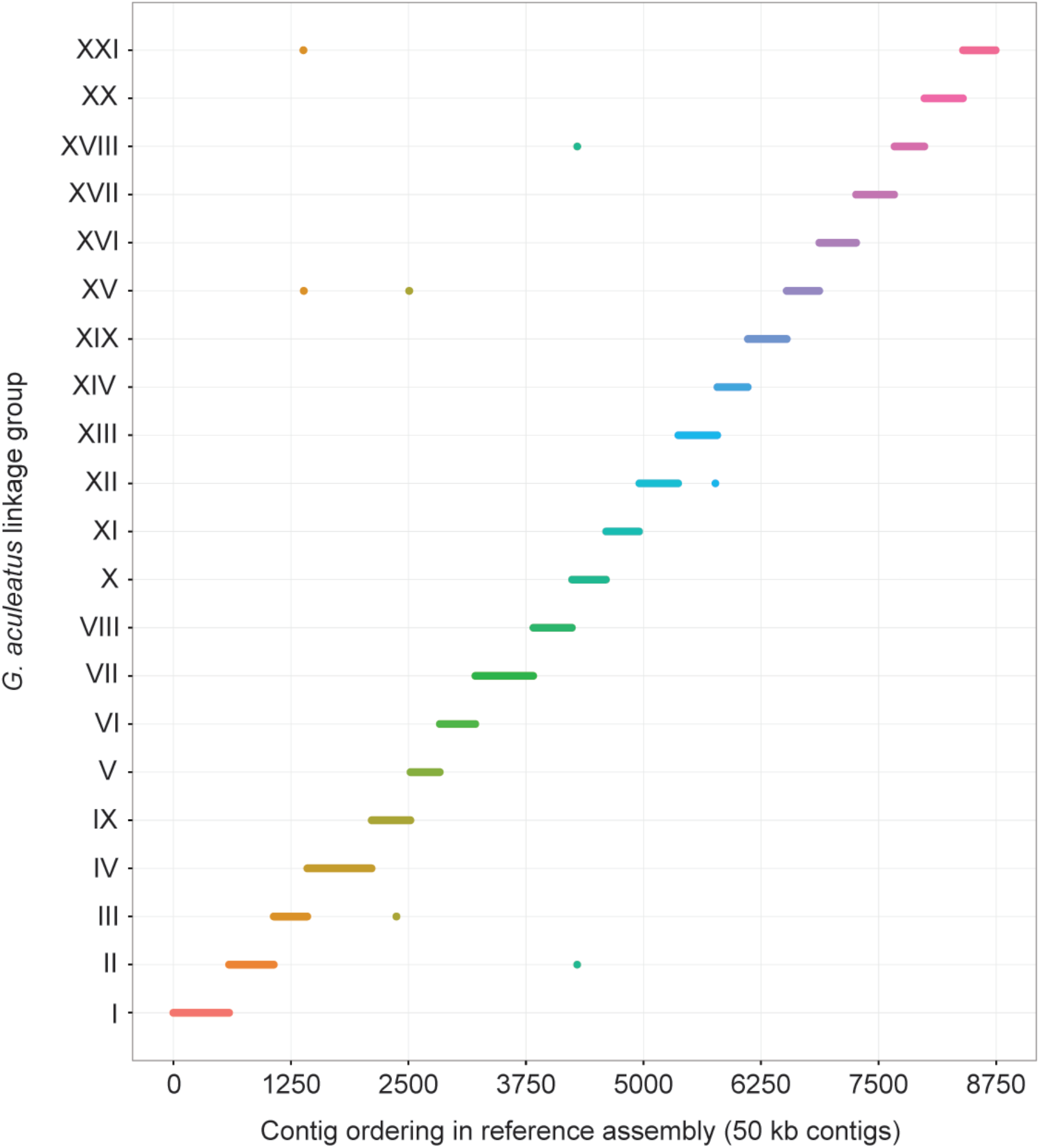
PGA clusters all linkage groups of the *G. aculeatus* genome. The revised *G. aculeatus* genome assembly (Glazer et al. 2015) was divided into contiguous 50 kb contigs and assembled using proximity-guided assembly (PGA). The *G. aculeatus* revised reference assembly contig order is preserved along the X-axis. PGA clustering was largely congruent with the revised *G. aculeatus* reference genome, as shown by each linkage group (LG) assembled as a contiguous segment. Seven contigs were assigned to different linkage groups in the PGA clustering than in the reference assembly. For example, a contig that was assigned to LG IX in the reference assembly is assigned to LG III in the PGA cluster (brown dot in the lower left part of the figure).

**Figure 2.**
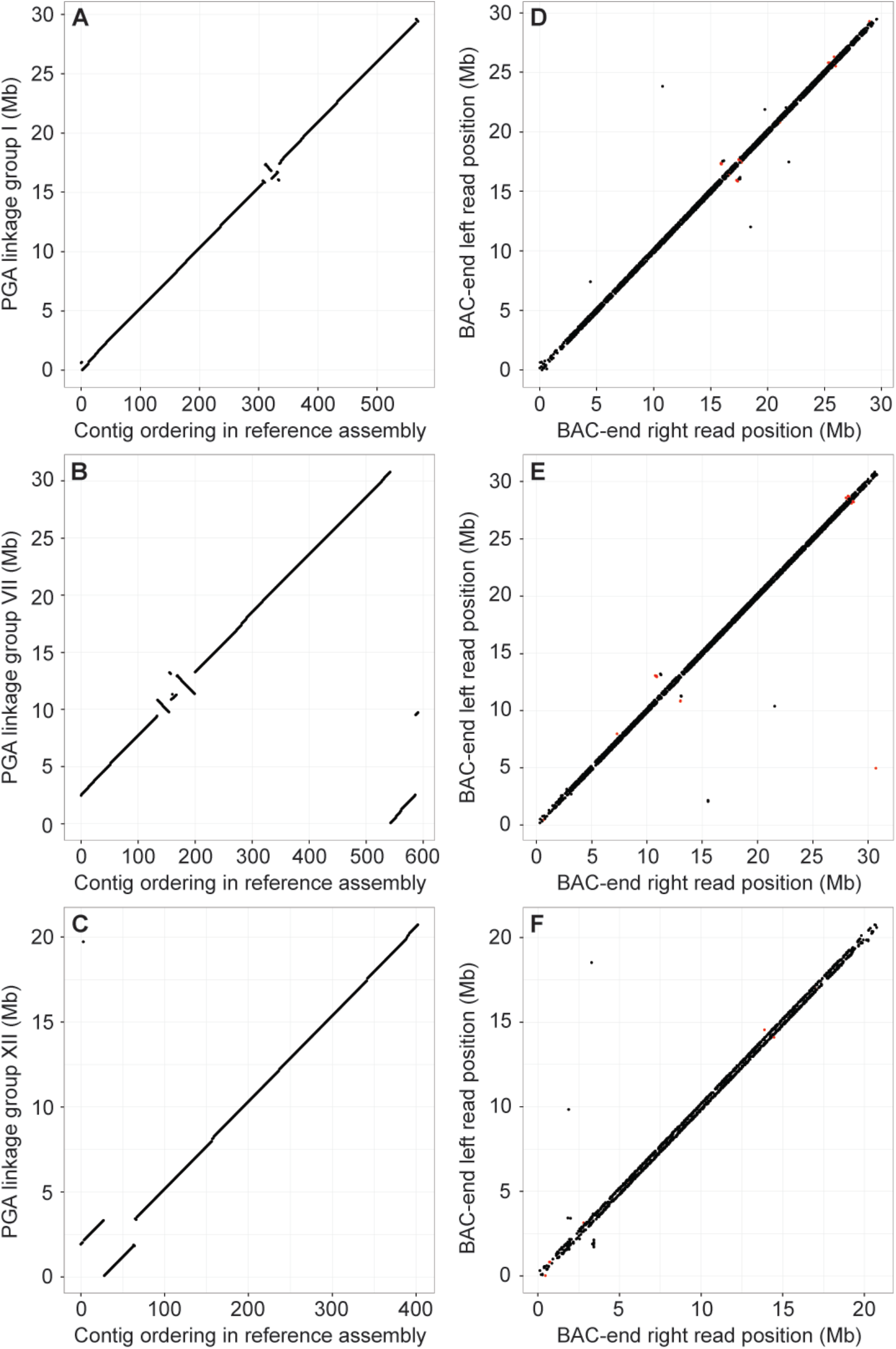
Misorderings within linkage groups are consistent with structural variation between populations of *G. aculeatus*. Misorderings in the PGA scaffolding within linkage groups (A-C) and alignment of BAC mate-pair sequences (D-F) relative to the *G. aculeatus* revised reference assembly (Glazer et al. 2015) are shown for three linkage groups that had significant overlap between the two data sets: (LG) I (A,D), VII (B,E), and XII (C,F). Alignment of BAC-end sequences from the same Paxton Lake benthic population used for the PGA scaffolding reveal structural variation around the breakpoints of the misordered PGA scaffolds. Left and right aligned mate-pairs from the BAC library largely follow a diagonal, reflecting the 148 kb average insert size of the library (Kingsley et al. 2004). Discordant read pairs fall off the diagonal. Black points indicate mate pairs that align in a forward/reverse orientation, reflecting a putative insertion relative to the reference genome assembly. Red points indicate mate pairs that align in a forward/forward or reverse/reverse orientation indicating a putative inversion relative to the reference genome assembly. The remaining linkage groups are shown in Figure S1.

We explored whether the incorrect orderings within linkage groups were due to errors in PGA scaffolding or structural variation using the BAC-end sequences available from a BAC library made from two Paxton Lake benthic males (Kingsley et al. 2004; Kingsley and Peichel 2007). We aligned the BAC-end sequences to the revised *G. aculeatus* genome assembly (Glazer et al. 2015) and scanned for discordant mate-pair alignments (i.e. mate-pairs that aligned in the same orientation or that aligned across genomic regions larger than 250 kb, which is larger than the 148 kb average insert size of the library). Such discordant alignments would indicate structural differences between the Paxton Lake benthic population and the Bear Paw Lake reference assembly population, or errors in the Bear Paw reference genome assembly. In many linkage groups, we found that discordant orderings between PGA scaffolding and the genome assembly were significantly more often associated with regions of the genome where BAC mate pairs aligned discordantly (Figure 2; Figure S1; Table 1). This indicates that at least some of the PGA scaffold misorderings may reflect true insertions, deletions, or inversions between populations of *G. aculeatus*, and/or errors in the reference assembly.

**Table 1.** Misorderings in the PGA scaffolding often overlap with BAC-ends that align discordantly to the reference genome.

| Linkage group | PGA Misorderings (N) | Discordant BAC ends (N) | Overlap (N) | P-value |
| --- | --- | --- | --- | --- |
| I | 6 | 104 | 5 (83%) | <0.001 |
| II | 4 | 32 | 1 (25%) | 0.133 |
| III | 2 | 10 | 0 (0%) | - |
| IV | 3 | 46 | 0 (0%) | - |
| V | 2 | 14 | 1 (50%) | 0.054 |
| VI | 2 | 16 | 2 (100%) | <0.001 |
| VII | 9 | 76 | 4 (44%) | 0.002 |
| VIII | 3 | 22 | 1 (33%) | 0.120 |
| IX | 4 | 30 | 0 (0%) | - |
| X | 6 | 102 | 5 (83%) | <0.001 |
| XI | 1 | 32 | 0 (0%) | - |
| XII | 4 | 56 | 2 (50%) | 0.039 |
| XIII | 2 | 52 | 1 (50%) | 0.104 |
| XIV | 3 | 6 | 0 (0%) | - |
| XV | 0 | 4 | - | - |
| XVI | 1 | 64 | 0 (0%) | - |
| XVII | 0 | 6 | - | - |
| XVIII | 0 | 20 | - | - |
| **XIX** | 4 | 70 | 2 (50%) | 0.179 |
| **XX** | 0 | 16 | - | - |
| **XXI** | 16 | 38 | 1 (6%) | 0.287 |
| **Total** | 72 | 816 | 25 (35%) | <0.001 |

### Placement of unassembled G. aculeatus contigs

In the most recent *G. aculeatus* genome assembly, 26.7 Mb of sequence (including Ns) still remains unanchored to linkage groups (Glazer et al. 2015). Here, we used the PGA scaffolding data to fill gaps and to assign these unassembled scaffolds to linkage groups. The assembled linkage groups and unassembled scaffolds of the *G. aculeatus* genome were split into their underlying contigs (from the assembled portion of the genome: 13,435 contigs, total length after removing Ns: 424.9 Mb, median contig length: 10,661 bp, N50=87,544 bp; from the unassembled portion of the genome: 3,499 contigs, total length after removing Ns: 21.7 Mb, median contig length: 3,076 bp, N50=10,954 bp) and reassembled using PGA. Including the previously unassembled contigs slightly reduced the accuracy during the clustering step when compared to the assembly that did not include these contigs (Figure 1). This clustering resulted in 11,927 contigs (88.78% of the previously assembled contig count) from the assembled portion of the genome clustered correctly by linkage group, 1,381 previously assembled contigs not clustered to linkage groups (10.28% of the previously assembled contig count), and 127 contigs assigned to the incorrect linkage group (0.94% of the previously assembled contig count) (Figure 3). Of the unassembled contigs, 2,015 contigs (57.59% of the total unassembled contig count) were clustered with linkage groups. During the ordering step, 1,604 were scaffolded by PGA (45.84% of the total unassembled contig count), but 216 contigs could not be assigned to a single linkage group and were not considered further. The remaining contigs were split into two groups based upon the confidence of their placement within a linkage group (see methods). 125 contigs from the unassembled scaffolds were unambiguously placed in gaps between sequential contigs in the revised genome assembly. This resulted in an additional 1.1 Mb of sequence (5.1% of the total unassembled length) scaffolded into the *G. aculeatus* genome assembly. The remaining 1,263 unassembled contigs (12.25 Mb, 56.4% of the total unassembled length) were mapped to regions of linkage groups (median range: 832.8 kb; max range: 33.6 Mb; min range: 8,958 bp), but could not be assigned to specific gaps in the genome assembly (Table 2).

**Figure 3.**
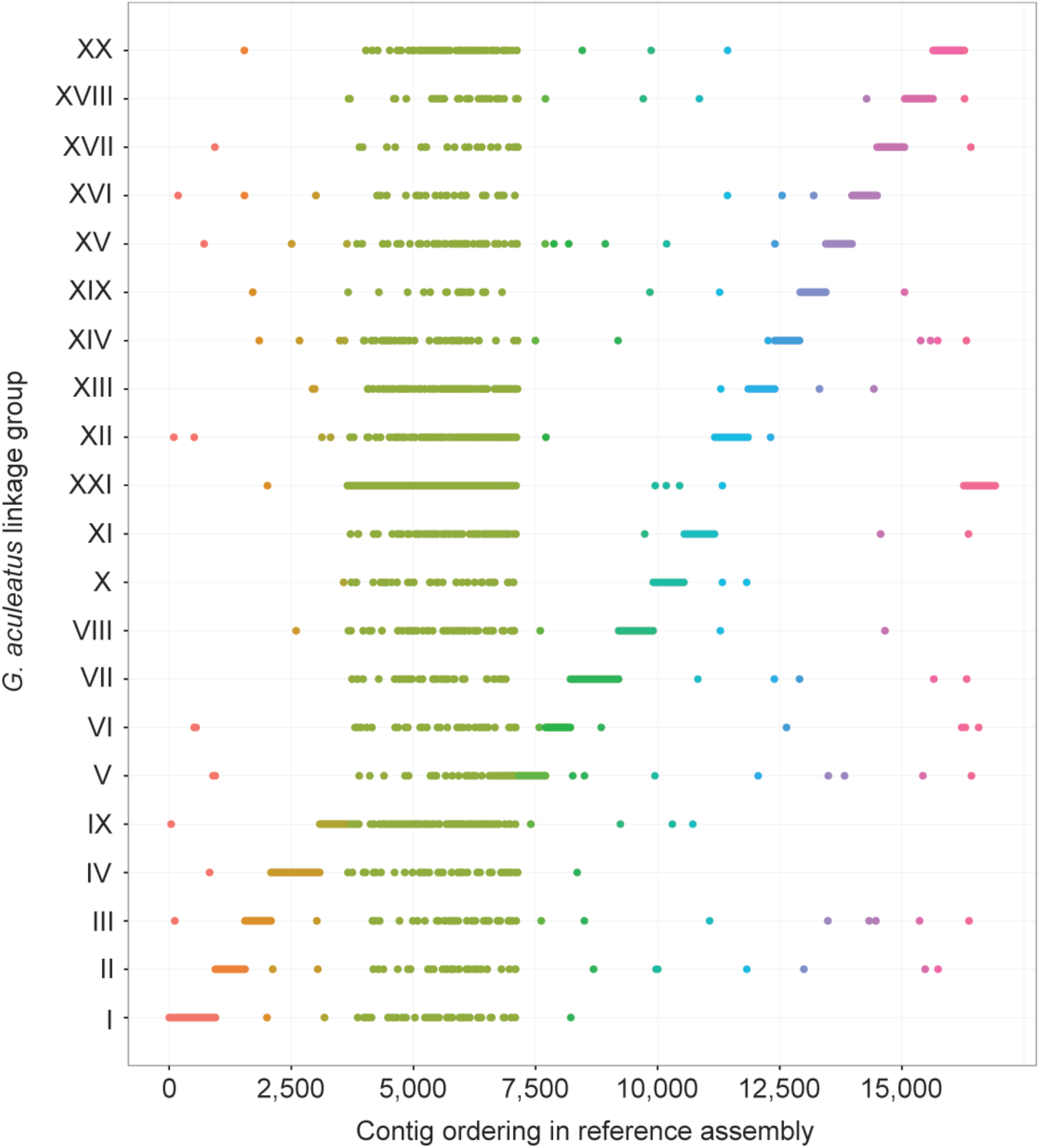
PGA clusters a large proportion of previously unassembled contigs to linkage groups. The revised *G. aculeatus* genome assembly (Glazer et al. 2015) was split into contigs at gaps and clustered with proximity-guided assembly (PGA) along with 21.7 Mb of contigs previously unassigned to linkage groups in the *G. aculeatus* genome. The *G. aculeatus* revised reference assembly contig order is preserved along the X-axis. The previously unassembled contigs (green vertical band) are distributed across linkage groups (LG) by the PGA clustering. PGA is less accurate clustering the *G. aculeatus* genome when the unassembled contigs are included, shown by an increased number of contigs being incorrectly assigned to different linkage groups (127 incorrectly assigned contigs, 0.94% of the previously assembled contig count).

**Table 2.** Distribution of contigs scaffolded by PGA that were previously unassembled in the *G. aculeatus* genome.

| Linkage group | Contigs placed into gaps (N) | Contigs placed into gaps (total length, bp) | Contigs narrowed to region (N) | Contigs narrowed to region (total length, bp) |
| --- | --- | --- | --- | --- |
| I | 10 | 93,296 | 45 | 433,601 |
| II | 6 | 48,648 | 11 | 135,728 |
| III | 1 | 8,144 | 23 | 216,619 |
| IV | 4 | 92,707 | 54 | 455,301 |
| V | 7 | 34,105 | 20 | 108,253 |
| VI | 1 | 8,060 | 20 | 191,189 |
| VII | 6 | 26,377 | 39 | 487,407 |
| VIII | 8 | 68,459 | 46 | 475,569 |
| IX | 31 | 291,837 | 109 | 909,866 |
| X | 1 | 1,635 | 34 | 411,527 |
| XI | 7 | 41,888 | 42 | 333,362 |
| XII | 6 | 46,023 | 124 | 965,250 |
| XIII | 6 | 49,046 | 81 | 719,633 |
| XIV | 1 | 8,315 | 51 | 527,141 |
| XV | 5 | 64,865 | 24 | 180,933 |
| XVI | 7 | 45,332 | 17 | 145,214 |
| XVII | 0 | - | 15 | 111,395 |
| XVIII | 0 | - | 30 | 398,982 |
| XIX | 4 | 33,136 | 10 | 66,498 |
| XX | 4 | 24,291 | 39 | 370,734 |
| XXI | 10 | 123,249 | 429 | 4,607,221 |
| **Total** | **125** | **1,109,413** | **1,263** | **12,251,423** |

To refine the chromosomal regions of the contigs not placed into specific gaps, we used long-distance mate pair information from the Paxton Lake benthic BAC-end sequences (Kingsley et al. 2004; Kingsley and Peichel 2007) to identify connections with contigs within the linkage group. We identified BACs where one end of a BAC insert aligned to an unscaffolded contig, while the other end aligned to a contig within the linkage group assigned by the PGA scaffolding. Identifying such linkage associations allowed us to narrow the location of many unscaffolded contigs to approximately 148 kb, the average insert size of the BAC library. Of the 1,263 unassembled contigs assigned to linkage groups by PGA scaffolding, 229 had alignments with BAC-ends. Of these contigs, 195 (2.9 Mb, 23.6% of the sequence length) had BAC-end alignments that matched the PGA linkage group associations (85.2%), confirming that the PGA scaffolding can accurately localize segments of the genome that are challenging to assemble through traditional methods. A revised genome assembly is provided as supplemental data, with 125 new contigs placed into gaps in the assembled genome and 1,263 of the unassembled contigs narrowed to linkage groups (available from Dryad Digital Repository).

### Scaffolding with BioNano Irys optical maps

We also used the BioNano Irys system (San Diego, CA) to generate optical maps of the Paxton Lake benthic population genome in order to verify the PGA scaffolding and to help refine location estimates of the unassembled contigs. The BioNano optical map was composed of 615 total contigs (N50=1.35 Mb) with a total map length of 569.7 Mb. We split the *G. aculeatus* revised genome assembly (Glazer et al. 2015) into 4,377 consecutive 100 kb bins (the minimum recommended contig length for BioNano assemblies) for rescaffolding with the BioNano optical map contigs. Using the default filtering parameters and alignment thresholds, the automated scaffolding pipeline (Shelton et al. 2015) was unable to join many contigs into scaffolds. The N50 remained at 100 kb, assembling 150 of the 4,377 contigs into 52 scaffolds. To improve scaffolding, we reduced the filtering thresholds of the scaffolding software by half. This increased the N50 of the assembly from 100 kb to 796 kb, incorporating 2,293 of the 4,377 contigs into 460 scaffolds; however, this also increased the number of misassembled scaffolds. 78 of the 460 scaffolds (19.0%) contained contigs from more than one linkage group. In addition to scaffolding, we aimed to use the BioNano optical maps to narrow the range estimates of the 1,263 unassembled contigs within their PGA assigned linkage groups, but this was not possible because only two of the 1,263 unassembled contigs were over the 100 kb minimum length required for scaffolding with BioNano optical maps.

## Discussion

Hi-C-based PGA was able to accurately re-scaffold the *G. aculeatus* revised genome assembly from 50 kb contigs into full linkage groups with internal ordering that closely matched the reference genome. Our results indicate PGA is highly effective at scaffolding relatively short contigs together into a contiguous assembly. Illumina short-read sequences are widely used to construct *de novo* genome assemblies in non-model organisms (reviewed in Ekblom and Wolf 2014)). However, short sequencing reads typically cannot span highly repetitive segments of genomes (Gordon et al. 2016; Treangen and Salzberg 2012). This limits the length of contigs that can be built from short-read technologies alone, often with contig N50 sizes of 10-50 kb (Ekblom and Wolf 2014). To assemble contigs into larger scaffolds, many genome assemblies incorporate long range information from a variety of sources, including BAC and fosmid libraries (Myers et al. 2000; Salzberg et al. 2012), jump libraries (Salzberg et al. 2012; Nagarajan and Pop 2013), optical mapping (Shelton et al. 2015; Zhang et al. 2012; Dong et al. 2013), genetic linkage maps (Fierst 2015), and single-molecule real-time sequencing (Bickhart et al. 2016; Gordon et al. 2016; Shi et al. 2016). However, application of these technologies can be limited by cost, the ability to perform crosses, and the availability of material. For example, optical mapping and BAC library construction require isolation of high molecular weight DNA (Shelton et al. 2015; Teague et al. 2010; Kingsley et al. 2004), which is not possible in many situations.

Some technologies, like BioNano Irys optical mapping, also require a minimum contig length (Shelton et al. 2015) for scaffolding that is not typically achievable with short-read Illumina sequencing alone. For example, even with 100 kb contigs, we were not able to re-scaffold the *G. aculeatus* reference genome to the completeness observed with PGA scaffolding. Our results therefore offer a promising example of constructing a nearly-complete genome assembly *de novo* using only short-read technologies paired with PGA scaffolding.

Several unmapped contigs from the *G. aculeatus* reference assembly were either placed into specific gaps or localized to regions within linkage groups in the re-scaffolded genome. Among linkage groups, there was an overabundance of unmapped contigs that were assigned to LG XXI. This linkage group harbors a 1.7 Mb inversion between marine and freshwater populations of sticklebacks (Jones et al. 2012), and consequently there is no recombination across this region in genetic crosses between marine and freshwater populations (Glazer et al. 2015; Wark et al. 2012). The original *G. aculeatus* reference genome assembly anchored scaffolds to linkage groups using a genetic map from a cross with the inversion segregating (Jones et al. 2012). The compressed genetic map and/or lack of markers in the region of the inversion may have prevented placement of many of the contigs, which were correctly assigned by the PGA scaffolding.

Within linkage groups, we identified several discordant orderings between the Hi-C assembly and the *G. aculeatus* reference genome. Although some of these are likely due to errors in the PGA scaffolding (from errors in the assembly algorithm or regional differences in chromatin interactions), many of the misorderings in the Hi-C assembly matched discordant mate-pair alignments in a BAC library from the same population of *G. aculeatus* used for the PGA scaffolding, suggesting structural variation among populations. Three population-specific inversions on LG I, LG XI, and LG XXI have previously been identified in sticklebacks (Jones et al. 2012). These inversions were not identified by the PGA scaffolding or by aligning the BAC-ends to the reference genome. However, these inversions are polymorphisms present between freshwater and marine populations and so would not be expected in our comparison between two freshwater populations (Paxton Lake benthic and Bear Paw Lake). Future work will focus on identifying the nature of the discordant orderings in the PGA assembly, and whether they reflect errors in the reference assembly or structural polymorphisms between the Paxton Lake benthic and Bear Paw Lake populations. However, the results presented here suggest that PGA scaffolding may also be a useful method to identify errors in reference assemblies or structural variation across genomes.

## Methods

### Tissue collection and Hi-C sequencing

The liver of a single adult male from the Paxton Lake benthic population (Texada Island, British Columbia) was dissected and flash frozen in liquid nitrogen. Tissue processing, chromatin isolation, library preparation, and sequencing were all performed by Phase Genomics (Seattle, WA). A total of 176,461,081 read-pairs were sequenced.

### PGA scaffolding

Scaffolding was conducted in two phases. First, to scaffold the entire *G. aculeatus* revised genome assembly (Glazer et al. 2015), the genome was divided into contiguous 50 kb contigs, excluding contigs that were not assigned to linkage groups and the mitochrondria sequence. Paired-end reads were aligned to the contigs, only retaining reads that aligned uniquely. Contigs were scaffolded using Proximity-Guided Assembly with an adapted version of the Lachesis method (Burton et al. 2013) and custom parameters developed by Phase Genomics. The second phase of scaffolding used PGA to assign the contigs that were previously not assigned to linkage groups to gaps in the reference genome. The reference assembly (excluding the mitochondria sequence) was split into contigs at gaps and Ns were removed, and all contigs were assembled with PGA. Contigs from the 21.7 Mb of previously unassembled sequence that were placed in the PGA assembly were divided into three groups, based on the level of certainty in their placement. If the contigs assembled before (contig A) and after (contig B) the previously unassembled contig were sequential (i.e. occurred in the expected order relative to the reference assembly), the previously unassembled contig was considered an accurate placement and was inserted in the gap between contig A and contig B. If contig A and contig B were from the same linkage group, but were not sequential, the previously unassembled contig could not be accurately placed in the gap and was instead assigned to the linkage group within a narrowed range of possible locations. If contig A and contig B were from different linkage groups or if contig A or contig B were missing (i.e. the previously unassembled contig only had linkage information on one end), the previously unassembled contig could not be placed and was not considered further (216 total unplaced contigs).

### BAC-end alignments

Sequenced BAC-ends from the CHORI-215 BAC library made from two Paxton Lake benthic males (Kingsley et al. 2004; Kingsley and Peichel 2007) were aligned to the unmasked *G. aculeatus* revised genome assembly (Glazer et al. 2015) using the BLAST-like alignment tool (BLAT) (Kent 2002). Alignments were only retained if at least 90% of the length of the sequenced BAC-end aligned to the genome. If a BAC-end aligned to multiple locations in the genome, the highest scoring alignment was kept according to the formula: alignment matches+alignment matches that are part of repeats – mismatches in alignment – number of gap openings in the query sequence – number of gap openings in the target sequence. Alignments were discarded if there were multiple alignments tied for the same highest alignment score. To retain the most stringent alignments, alignments were also not considered if they contained gaps totaling more than 10% of the alignment length.

### Overlap between misorderings in the PGA assembly and BAC-ends that aligned discordantly

BAC-ends from the same insert were considered to align discordantly if they were separated by greater than 250 kb in the genome (the average insert size of the library is 148 kb), regardless of orientation of the BAC-ends. Alignments were only considered if both BAC-ends aligned to the same linkage group, reflecting intrachromosomal rearrangements. BAC-ends that aligned in a forward/reverse orientation and were separated by over 250 kb in the genome were considered putative insertions in the Paxton Lake benthic population or deletions in the reference assembly. BAC-ends that aligned in a forward/forward or reverse/reverse orientation and were separated by over 250 kb in the genome were considered putative inversions. A contig was considered misordered in the 50-kb PGA reassembly of the *G. aculeatus* genome if the previous contig in the scaffold was located further than 250 kb away from the coordinates in the reference genome assembly (Glazer et al. 2015). Overlap was scored if the position of a discordantly aligned BAC-end fell within the 50 kb contig that was misordered in the PGA assembly. Permutations were conducted for each linkage group separately and for the total genome to test for significance. Random subsets of BAC ends were drawn from each linkage group equal to the number of discordant BAC-end alignments. Overlap was scored between the misordered PGA contigs and the random subsets of BAC ends. The p-value reflects how often the same number of overlaps is recovered among a set of 10,000 random permutations.

### BioNano scaffolding

High molecular weight DNA was isolated from the blood of a single adult male from the Paxton Lake benthic population (Texada Island, British Columbia) following protocols outlined in (Kingsley et al. 2004). This male was not the same individual used for Hi-C, nor for creation of the BAC libraries. Irys optical mapping (BioNano Genomics, San Diego, CA) was performed at Kansas State University. DNA was nicked with the BspQI restriction enzyme, which cuts at a frequency of 15.8 sites per 100 kb across the *G. aculeatus* genome, which is around the ideal cutting frequency of 10-15 sites / 100 kb for optical mapping with the BioNano Irys System (Shelton et al. 2015). DNA was labeled with fluorescent nucleotides and repaired according to BioNano protocols. DNA was imaged on the BioNano Irys System using two IrysChips. BioNano molecules were filtered to only include segments that were at least 150 kb and contained at least eight labels. The P-value threshold for the BioNano assembler was set to a minimum of 2.2×10^−9^. Molecule stretch was adjusted using AssembleIrysCluster.pl (v. 1.6.1) (Shelton et al. 2015). To assess the effectiveness of BioNano optical maps in rescaffolding the *G. aculeatus* genome, the revised genome assembly was split into contiguous 100 kb contigs (the minimum recommended size for BioNano assembly). The split genome was digested with BspQI into *in silico* CMAP files using fa2cmap_multi.pl (BioNano) and iteratively scaffolded with the BioNano optical maps using sewing_machine.pl (v. 1.0.6) (Shelton et al. 2015). Two different filtering options were used: the default filters (--f_con 20, --f_algn 40, --s_con 15, --s_algn 90) and relaxed filters set at half of the default thresholds (--f_con 10, --f_algn 20, --s_con 7.5, --s_algn 45). For both sets of filters, the default alignment parameters were used (-FP 0.8, -FN 0.08 –sf 0.20 –sd 0.10).

## Acknowledgements

We thank Chris Amemiya for isolating high molecular weight DNA for use in optical mapping, and Susan Brown and the Kansas State University Bioinformatics Center for performing the optical mapping. This work was supported by an Evolutionary, Ecological, or Conservation Genomics Research award from the American Genetic Association (MAW), the Office of the Vice President of Research at the University of Georgia (MAW), NIH grant R01 GM116853 (CLP), and the Fred Hutchinson Cancer Research Center Division of Basic Sciences (CLP).

## Data Access

Hi-C sequences are deposited in the NCBI SRA database (SRP081031). The revised genome assembly and the XMAP optical map files will be available on the Dryad Digital Repository.

## Author Contributions

CLP and MAW conceived and designed experiments, CLP, STS, IL and MAW analyzed data and interpreted results, and CLP and MAW wrote the manuscript with input from STS and IL.

